# A can of worms: estimating the global number of earthworm species

**DOI:** 10.1101/2024.09.08.611896

**Authors:** Thibaud Decaëns, George G. Brown, Erin K. Cameron, Csaba Csuzdi, Nico Eisenhauer, Sylvain Gérard, Arnaud Goulpeau, Mickaël Hedde, Samuel W. James, Emmanuel Lapied, Marie-Eugénie Maggia, Daniel F. Marchán, Jérôme Mathieu, Helen R. P. Phillips, Eric Marcon

## Abstract

Estimating the overall species number for a given taxon is a central issue in ecology and conservation biology. This is especially topical for soil organisms, which comprise most known species but whose taxonomy remains largely understudied. Here, we estimated the global number of earthworm species based on the Joppa approach, which models taxonomic effort over time to estimate the total number of known and as yet unknown species in a given taxa. Our Bayesian estimation of the Joppa model suggests a global diversity of the order of 30,000 species, suggesting that the 5,679 earthworm taxa already described only represent around 20% of the actual global species diversity. However, the uncertainty around this estimate is considerable due to severe undersampling and as the model cannot unambiguously decide whether we are describing few species because of a small pool of as yet unknown species, or because of a lack of taxonomic efficiency. Considering the current rate of new species description, we calculate that it would take at least 120 years to describe all the earthworm species existing on Earth, and we discuss thedifferent strategies that should be developed to facilitate and accelerate the discovery and naming of species new to science.

## Introduction

Estimating the total number of species of a given taxon, whether on a global scale or for a given region or ecosystem, is a question of great practical and symbolic importance that has long captivated and challenged scientists (May, 1992; Stork, 1993; Pimm, 2001; Dirzo & Raven, 2003). It represents an indispensable step towards a comprehensive description of the World’s biodiversity and is also necessary for the definition of efficient conservation strategies (Joppa et al., 2011; Caley et al., 2014). In this context, soil biota represents one of the most diversified and least studied compartments in terrestrial ecosystems. They are among the most diverse, comprising up to 59% of currently known living species (Decaëns et al., 2006; Anthony et al., 2023). They are involved in regulating a large number of ecological processes that are essential to the provision of key ecosystem services for human populations (Brussaard et al., 1997; Wardle et al., 2004; Bardgett & van der Putten, 2014). This has been particularly well documented for emblematic groups of soil ecosystem engineers, such as earthworms (Lavelle et al., 2006, 2016). Paradoxically, even these well-studied groups still suffer from a glaring lack of basic studies on their biodiversity patterns (Cameron et al., 2018; Phillips et al., 2019, 2021), and questions as fundamental as “how many species are there?” still remain largely unanswered (Decaëns, 2010). This knowledge gap is of particular concern, as it limits our capacity to characterize the dynamics of soil biodiversity in the face of ongoing global change and to assess its implications for ecosystem functioning.

Earthworms are an emblematic illustration of this paradox. Some 5,700 earthworm species have been described to date (Misirlioğlu et al., 2023), but the proportion of the actual diversity of these organisms that this figure represents is still poorly documented. Estimates vary, depending on the author, from a minimum of 7,000 to some 20,000 species (Reynolds, 1994; Lavelle et al., 2006; Blakemore, 2009; Anthony et al., 2023), but most of these figures are based on expert opinion, and no mathematical formalisation has yet been used to obtain a consolidated prediction of the number of as-yet undiscovered species. Some studies have also highlighted the existence of potentially high and as yet largely undescribed regional diversity throughout the tropics (Phillips et al., 2019; Goulpeau et al., 2024), which could challenge these previously proposed estimates of global species number.

Various methods can be used to estimate the number of heretofore unknown species within a given taxon. One popular approach is to use rarefaction and extrapolation curves, combined with non-parametric estimators, to predict the taxonomic deficit and the number of species at different spatial scales (Chao, 1984; Colwell et al., 2004; Chao et al., 2005; Chao & Jost, 2012; Cazzolla-Gatti et al., 2022). However, to be effective, this approach requires a sufficiently dense dataset on a global scale, which is lacking for many soil organisms such as earthworms. Another method consists of using Arrhenius’ power law to estimate the number of species for large areas on the basis of a known number of species in a smaller area (Arrhenius, 1921; Krishnamani et al., 2004; Marcon et al., 2024). The procedure involves estimating a power parameter (*z*) based on the negative linear relationship between the log of compositional similarity and the log of geographical distance between small equivalent areas. It therefore requires a network of plots located at very different distances, which must be supplemented by a continuous inventory over an area corresponding to the smallest scale of the power law. Apart from the fact that this type of data is rare, log transformation is impossible for pairs of plots with a compositional similarity of 0 (i.e. no species in common), which is common in the case of earthworms, particularly in the tropics (Goulpeau et al., 2024). Finally, Joppa *et al*. (2011) propose to use global species checklists to model the rate at which taxonomists describe new species as a function of time, and to estimate the number of species that will have been reached when this rate will drop to zero due to the exhaustion of the pool of as yet unknown species. This approach has the advantage of not being dependent on the existence of comprehensive field datasets, but requires the establishment of reliable species checklists that can be challenged by taxonomic issues.

In this work, we propose a global estimate of the number of earthworm species based on the method of Joppa *et al*. (2011) and the most recent global checklist of earthworms (Brown et al., 2023; Misirlioğlu et al., 2023). In parallel, we analysed three datasets obtained from field inventories for regions equivalent in surface area in tropical and temperate regions (Bouché, 1972; Goulpeau et al., 2024; Gérard et al., 2024). We used these data to quantify the extent of the taxonomic deficit and estimate the actual number of species for each region. We then used these regional estimates to discuss the range of results obtained from the Joppa method.

## Material and methods

### Estimating the number of species from the global checklist

For the Joppa approach, we used the checklist recently published by Misirlioğlu *et al*. (2023) and Brown *et al*. (2023). This list includes 5,738 species/subspecies (5,406 species and 332 unique subspecies, i.e. not counting nomino-typical subspecies) belonging to 23 families (including one non-crassiclitellate family: Moniligastridae). We chose to consider subspecies as taxa in their own right in the analysis, because the choice of whether to consider them as infraspecific divisions or to elevate them to species rank is a matter of taxonomists’ points of view. For the sake of simplicity, we referred to all the taxa in the list as “species”. After deleting certain duplicates in the table, we obtained a consolidated list of 5,752 taxa names published between 1758 and 2023, containing for each of them the authors and the year of description. For each 5-year period, we then counted the number of unique species described and the total number of authors (i.e. all the co-authors of the species, after standardising the names of authors with accents and taking account of homonyms, wherever possible).

Joppa *et al*. (2011) expected the number of species (*s*_*i*_) described in a given interval (*Y*_*i*_ = *i*, numbered sequentially from 1) to depend on the taxonomic effort, i.e. the number of taxonomists (*T*_*i*_) actively describing species during that period, and the unknown number of species remaining to be described, that is the total number of species (*s*_*T*_) minus those already discovered 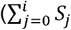 where *s*_*0*_ is the number of known species before *Y*_*1*_). Over time, taxonomists were expected to become more efficient, i.e. the discovery rate of unknown species per taxonomist was assumed to increase linearly. The original model was thus

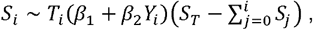

where *β*_*l*_ and *β*_2_ are unknown parameters. The model was log-transformed and estimated with a normal error of variance *σ*^*2*^. The parameters to be estimated were thus *β*_*l*_, *β*_2_, *s*_*T*_ and *α*, the total number of species *s*_*T*_ being our main target.

The increase in efficiency accounted for in the Joppa model is not obvious as earthworms are concerned. Methodological advancements, such as the development of integrative taxonomy, are likely offset by the increased difficulty of sampling in remote areas, namely tropical rainforests. Moreover, contemporary scientists rarely focus exclusively on taxonomy, a discipline whose publications are generally undervalued, leaving them with less time to describe new species. Consequently, we studied an alternative model with no efficiency progress (hereafter referred to as the “no-progress” model), where *β*_2_ equals zero.

We chose Bayesian inference to obtain the distribution of the estimates of the parameters which revealed to be as important as the most probable values. We chose non-informative priors, uniformly distributed between 0 and numerically possible values *β*_*l*_, *β*_2_ and a. According to expert opinion, the prior of s_T_ was forced between 6,000, i.e. the number of already discovered species, and 100,000. The parameter space was explored by a Monte-Carlo Markov Chain with 10^7^ steps, 10% considered as burn-in. The analysis began with time period 1822-1827, with a previously known number of species (*s*_*0*_) equal to 1 (i.e. *Lumbricus terrestris* Linnaeus, 1758). The last period is 2018-2022; species published in 2023 and 2024 were excluded from the analysis.

### Estimating regional diversity and the taxonomic deficit

Compared to the checklist of angiosperms used by Joppa *et al*. (2011), that of earthworms is very short: data is scarce. We endeavoured to supplement the global approach with three more exhaustive regional studies.

For the analyses of regional diversity, we used the data of Goulpeau *et al*. (2022, 2024) in French Guiana (supplemented by a small number of as yet unpublished data), and, for Southern and Northern France, those of Bouché (1972) updated for taxonomy by Gérard *et al*. (2024). The choice of these three regions allows us to consider contrasting situations in terms of earthworm diversity. French Guiana is considered to be a diversity hotspot, with a high proportion of species with restricted distribution and a large taxonomic deficit (Decaëns et al., 2016, 2024; Maggia et al., 2021; Goulpeau et al., 2024). By comparison, mainland France has a lower *a priori* species diversity, but the South is nevertheless characterised by a high proportion of endemic species, some of which continue to be discovered regularly, whereas the North is characterised by great compositional uniformity (Bouché, 1972; Mathieu & Davies, 2014; Gérard et al., 2023; Marchán et al., 2023).

Three 300 × 400 km^2^ zones, each subdivided into twelve 100 × 100 km^2^ cells, were delimited in each region. In each 100 × 100 km^2^ cell, we counted the species present in order to obtain three occurrence tables comprising 12 rows and as many columns as species present. In French Guiana, we used the operational taxonomic units (OTUs) proposed by Goulpeau *et al*. (2022) as a substitute for species in order to overcome the problem posed by the taxonomic deficit. For each region separately, we plotted incidence-based rarefaction and extrapolation curves in order to represent the accumulation of species richness as a function of the number of 100 × 100 km^2^ cells sampled in the study areas. The 95% confidence intervals of the interpolated and extrapolated richness values were obtained by a bootstrap procedure with 200 replications. We also calculated the non-parametric Chao 2 index, which estimates the lower limit of expected asymptotic species richness (Gotelli & Chao, 2013). The taxonomic deficit, i.e. the proportion of undiscovered species, was calculated using the formula:

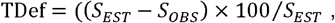

with *TDef* the taxonomic deficit, *s*_*OBS*_ the number of species currently known for a given region, and *s*_*EST*_ the richness estimated by the Chao 2 index.

For French Guiana, the taxonomic deficit was calculated by first considering *s*_*OBS*_ as the set of OTUs delimited by Goulpeau *et al*. (2022), and then by restricting *s*_*OBS*_ to the number of species formally described to date (i.e. 55 species taking into account the 18 new descriptions by Decaëns *et al*. (2024). These analyses were performed using the *‘iNEXT’* package for R (Hsieh et al., 2019; R Core Team, 2024).

## Results

During the period from 1822 to 2022, the number of species described per 5-year period showed two peaks: the first from the end of the 19th to the middle of the 20th century, the other from the 1970s to the present day (Fig. 1A). Over the same period, the total number of authors of original species descriptions has increased exponentially (i.e. roughly linearly on a logarithmic scale; Fig. 1A). The ratio of numbers of species described per taxonomist thus shows a very marked peak in the first half of the 20th century, a period during which it generally exceeds 15 species described per taxonomist per time period, whereas everywhere else it is generally below 10 species per taxonomist (Fig. 1B).

**Figure 1.**
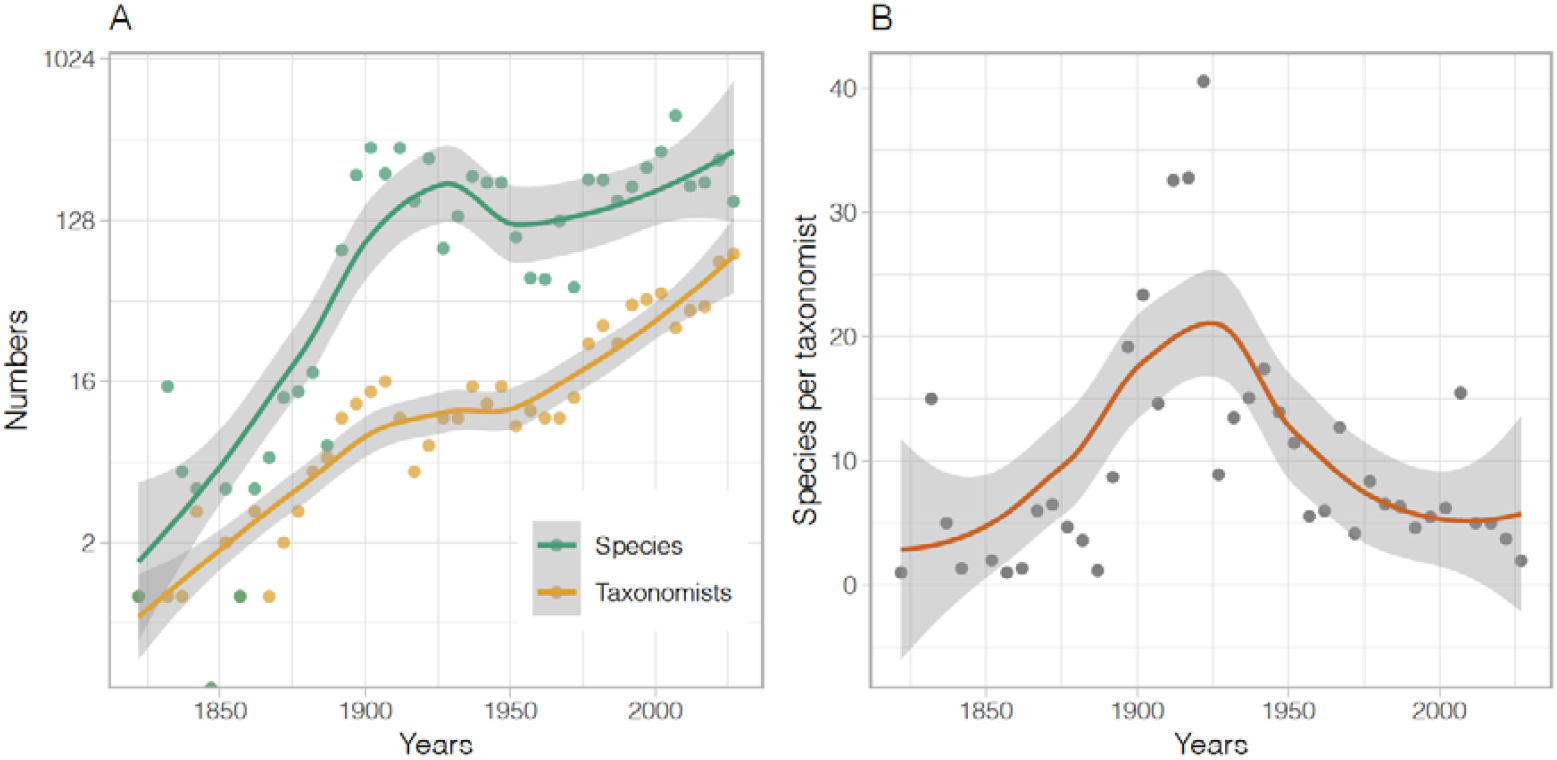
Fluctuations over time, for each 5-year period from 1822 to 2017, in the numbers of described earthworm species and taxonomists (A), and in the number of species described per taxonomist (B). The loess models fitted to the data are shown in colour, with the grey area corresponding to the 95% confidence interval.

The Joppa model yields the median posterior ST equal to 6,814 species, with a 95% higher posterior density interval (HPDI) between 6,000 and 11,148. The no-progress model leads to quite different estimations: 26,283 species (HDPI between 6,003 and 87,425). Posterior probabilities are shown in Fig. 2.

**Figure 2.**
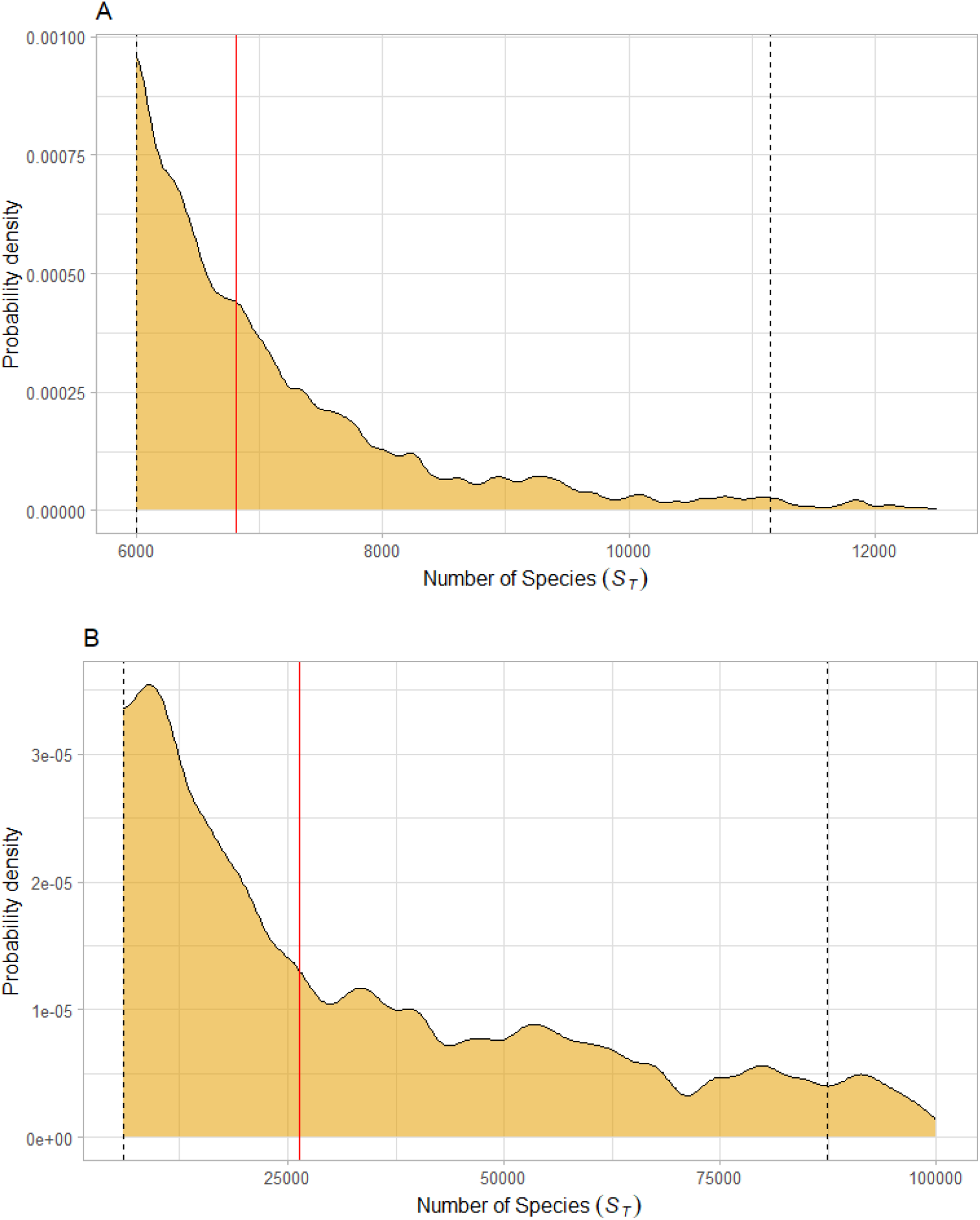
Results of Bayesian estimation of the global number of earthworm species according to the original Joppa model (A) and the no-progress model assuming the taxonomists’ efficiency constant (B). The diagram shows the probability density of the estimated ST value. The red line represents the median and the dashed lines are the limits of the highest posterior density interval.

The rarefaction curves obtained for the three study regions show a decrease in beta diversity from tropical to temperate study regions (Fig. 3). In French Guiana, the slope of the curve is steep, showing that the asymptote is far from being reached, whereas it has been or is about to be reached in the two study regions in temperate France. The diversity metrics logically follow the same trend: the cumulative number of species observed is 305 in French Guiana, 66 in Southern France, and 31 in Northern France, whereas the Chao 2 index predicts a diversity of 1,680 species in French Guiana, 99 in Southern France, and 36 in Northern France (Fig. 4A, 4B). Consequently, the taxonomic deficit is of the order of 80% in French Guiana, 33% in the south of France, and 16% in the north of France (Fig. 4C). In French Guiana, this 80% taxonomic deficit was obtained by considering as observed richness all of the OTUs mentioned in Goulpeau et al. (2022), most of which are new to science and have not yet been named. If we consider only the 55 named species actually known from French Guiana (Fig. 4A), the estimated taxonomic deficit rises to 97% (Fig. 4C).

**Figure 3.**
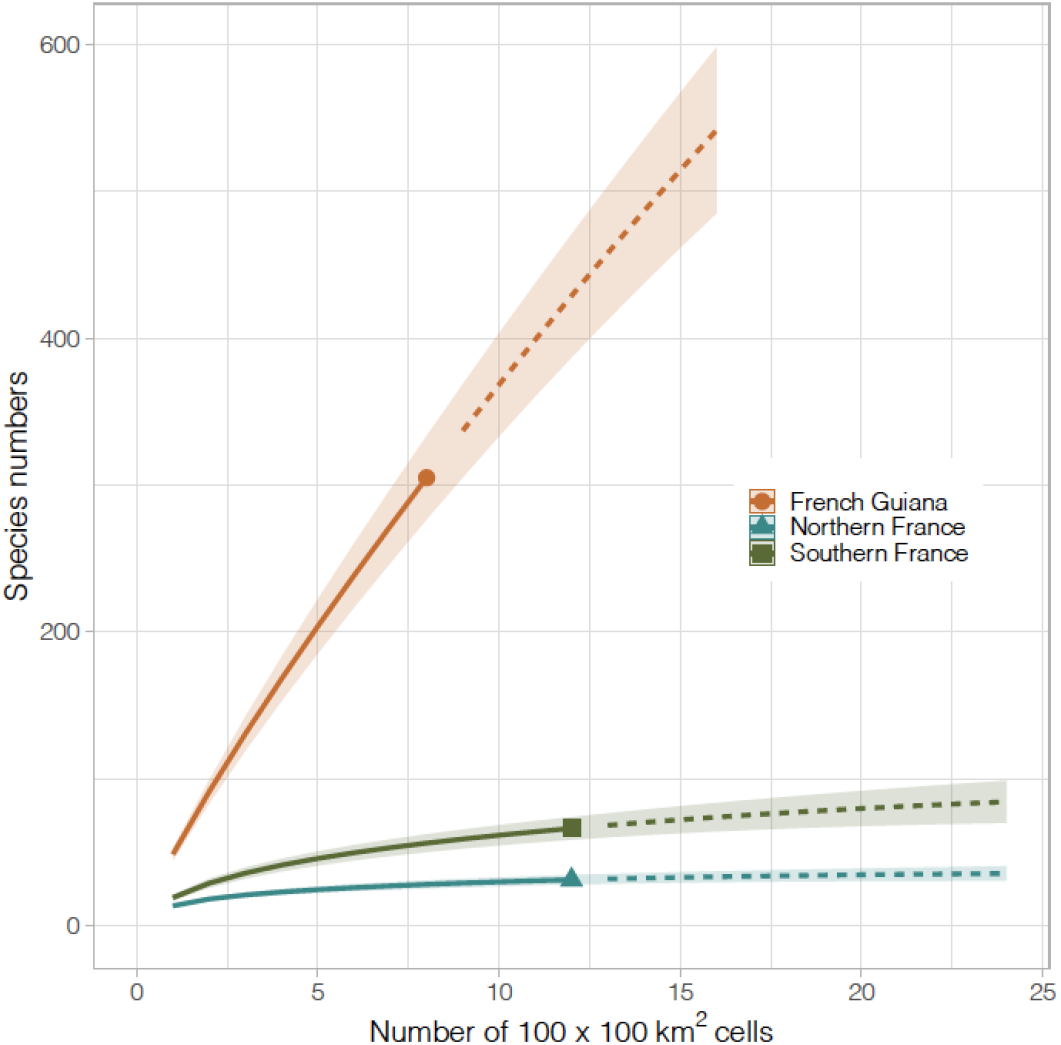
Incidence-based rarefaction and extrapolation curves showing how earthworm species richness increases as a function of sampling effort (number of 100 × 100 km2 windows) in the three regions studied. Solid lines represent rarefaction curves (interpolated values); dashed lines represent extrapolation curves; shaded areas represent 95% confidence intervals obtained with a bootstrap of 200 replications.

**Figure 4.**
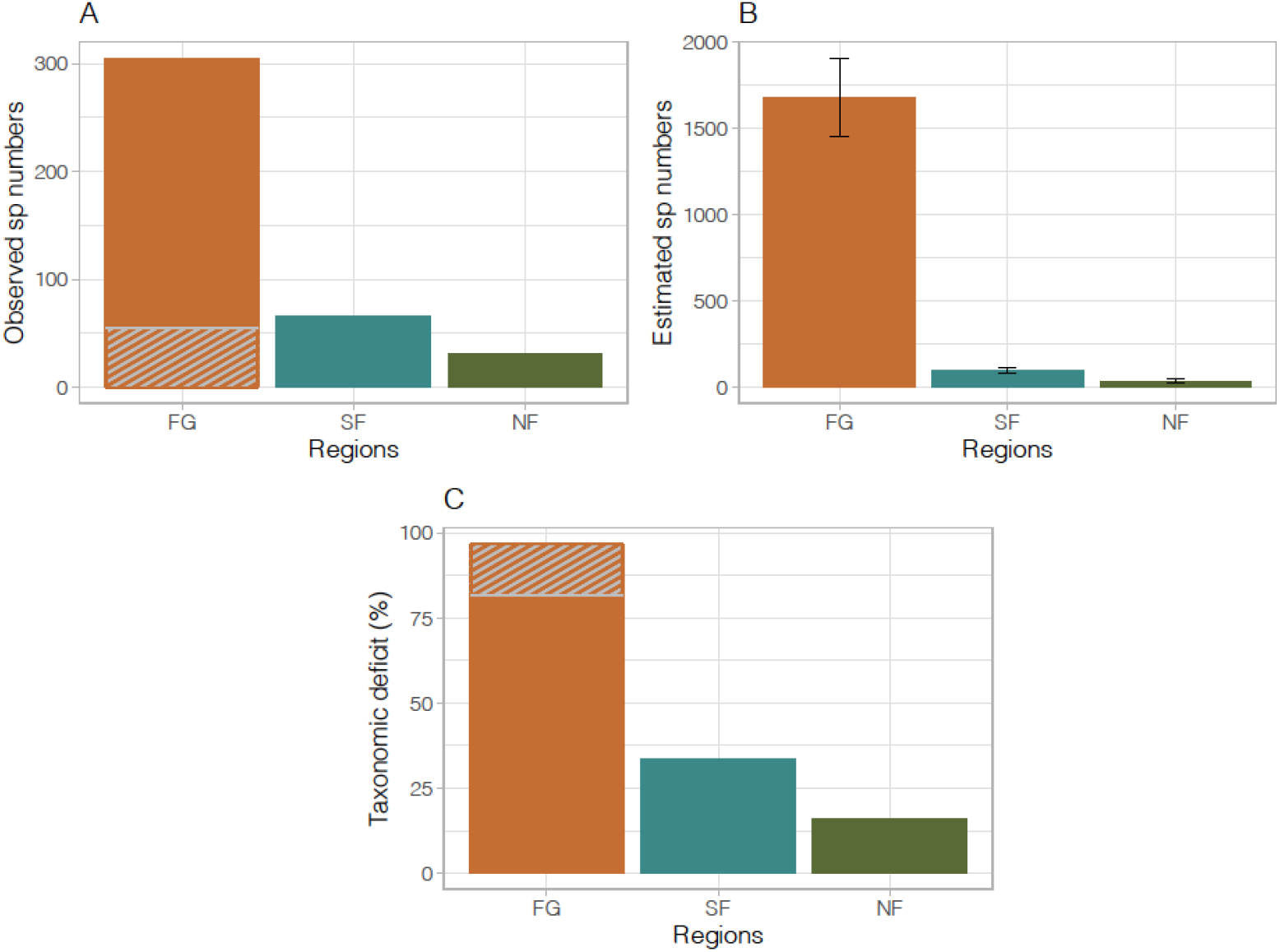
Analysis of earthworm diversity in the three study regions: A) cumulative number of observed species, B) estimated number of species (Chao 2 index) and C) taxonomic deficit (i.e. the proportion of estimated diversity that has not been observed). In A, the grey hatched part corresponds to species from French Guiana that have already been named, while the non-hatched part of the bar corresponds to species that have been discovered but not yet described; in B, the error bars represent the 95% confidence interval obtained using a bootstrap with 200 replications; in C, the hatched part corresponds to the inflation of the taxonomic deficit in French Guiana as a result of taking into account only species that have been already named, the non-hatched part of the bar corresponding to the deficit calculated by also taking into account species that have been discovered but not yet described.

## Discussion

Earthworm taxonomy experienced a golden age from the end of the 19th century to the middle of the 20th century, followed by a revival from the end of the 1970s. The first of these two periods can be credited to the great explorations that enabled zoologists of the time to explore the world and describe its biological diversity (Csuzdi & Szlávecz, 2016). It was also made possible by the exceptional productivity of the few taxonomists of the time, who were able to describe dozens of species in a short time. By comparison, the contemporary period is characterised by a larger community of taxonomists, but with comparatively lower output. Overall, we do not observe the monotonic decline in the number of species described per taxonomist as described for plants and other animal taxa (Pimm et al., 2010; Joppa et al., 2011; Fisher et al., 2018), which is assumed to reflect a depletion of the pool of new species to be discovered. Rather, with the exception of the notable peak in the first half of the 20th century, the rate of description of new earthworm species remained modest and constant over time, at around 5 species per taxonomist per 5-year period. Even in the early 2000s, we did not observe any clear effect from the development of molecular tools such as DNA barcoding and metabarcoding (sensu Hebert et al., 2003; Taberlet et al., 2012). This may seem surprising, given that their usefulness in the field of earthworm taxonomy has been widely acknowledged (Rougerie et al., 2009; Decaëns et al., 2013; Marchán et al., 2022), and their advent has been touted as the possible catalyst of a new golden age of taxonomy (Hebert & Gregory, 2005; Hajibabaei, 2012; Pennisi, 2019). Over the last few decades, the increase in the number of new earthworm species described per time period seems to be simply correlated with the number of active taxonomists.

Our estimates of the Joppa and the no-progress model tell us two different stories. Assuming a constant progress of the efficiency of taxonomists, the global number of species would be below 7,000, i.e. around 1,000 species remaining to be described. On the other hand, assuming constant efficiency, the global number of species would rather be around 30,000, i.e. the 5,679 species of our checklist are only the beginning of the story.

In both cases, the confidence intervals around our median estimates are considerable, motivating us not to provide a single significant figure in our estimates. This can be explained by two main causes of uncertainty. First, we lack data: more publications of new species would have allowed smaller, i.e. more, time intervals, thus more statistical power. Also, the Joppa model does not allow to completely disentangle the effects of the efficiency and the number of species: in the no-progress model, the estimates are highly correlated (Pearson correlation of the posteriors: -0,73), and the correlation is -0,55 between *β*_2_ and *s*_*T*_ in the original model. This issue could not be detected by Joppa et al. (2011) who relied on a frequentist estimation of the most likely number of species.

The analysis of regional diversity in French Guiana points in the direction of a high number of species: we estimate that the taxonomic deficit there is of the order of 97%, if only currently named species are considered in the calculation. An unknown thousand species may well all be there: a small area (actually around 1%) of Amazonia may thus be enough to find the species missing from the Joppa model. If we consider that around half of the earthworm species currently known are tropical (i.e. around 2,800 species), the generalisation of such a taxonomic deficit would lead us to estimate that more than 90,000 species could exist throughout the tropics. Of course, such a generalisation is highly debatable, but it does provide an argument for considering that the global number of unknown earthworm species may not be only a few thousands.

As expected, we calculate a lower taxonomic deficit in temperate regions, which have been better studied and host a more modest diversity. However, our results indicate that this deficit can still be significant—ranging from 15 to 30% in mainland France—suggesting that even in temperate regions, a substantial number of species remains undiscovered. The South of France, known for its high diversity and richness in endemic earthworms, exemplifies this, as evidenced by the recent descriptions of several new species, including some of large body size (Marchán et al., 2023). The persistence of a significant taxonomic deficit in Northern France is more unexpected. It may reflect a gap in our understanding of species distribution in the region (i.e. the Wallacean shortfall according to Lomolino, 2004), or the persistence of taxonomic confusion that can conceal the presence of certain species. A striking example of this is the discovery of the existence of two cryptic species that were mistakenly considered synonymous under the name L. terrestris (James et al., 2010).

These considerations allow us to propose a conservative estimate of the global number of earthworm species of the order of 30,000 species. This is above the upper range of commonly accepted predictions, i.e. from 10,000 to 20,000 species (Reynolds, 1994; Decaëns et al., 2006; Blakemore, 2009; Anthony et al., 2023). Our expert opinion is that there is still a large pool of undiscovered species, but that we lack the efficiency to discover them or describe them once they have been discovered. First, a low rate of new species description per taxonomist, compared with those active at the beginning of the 20th century, is expected due the fact that taxonomy is no longer considered a ‘competitive’ discipline in modern science (Wilson, 2004; Agnarsson & Kuntner, 2007). The pernicious effect of this situation is that contemporary authors are rarely pure taxonomists, as they are encouraged by the academic system to devote most of their research time to more ‘valuable’ areas of research, such as evolution or ecology. Added to this is the fact that taxonomists generally lack the time and financial resources for field sampling, particularly in the most diverse and yet least well-studied tropical regions (Guerra et al., 2020). The field scientists among us all have in mind regions that they know contain a diversity of as yet unknown species, but which they do not have the time or the resources to explore. Similarly, taxonomists have a multitude of species in their cupboards waiting to be described, but which they do not have the time to process.

Overall, our results suggest that at best we have only described one fifth of the global number of earthworm species. If we consider the current community of taxonomists (an average of 45 active taxonomists per 5-year period over the last 20 years), and the number of species they have managed to describe (an average of 4.5 species per taxonomist per time step), we calculate that it would take at least 120 years to describe all the species existing on Earth. Obviously, such an objective is certainly unattainable within a reasonable timeframe, but we can nonetheless think about the strategies that could be developed to reduce the magnitude of the taxonomic deficit. This will require maintaining a major taxonomic effort, through the training of young taxonomists, but also by raising the profile of taxonomic work in the scientific corpus. Agnarsson & Kuntner (2007) have proposed that original descriptions, and their underlying taxonomic hypotheses, should be formally cited in articles that refer to them – which is not systematically the case at present. This would allow the impact metrics of taxonomic journals to improve while more accurately reflecting their real scientific impact, thus encouraging scientists to invest more in taxonomy. We will also need to improve taxonomic efficiency, first by increasing our ability to discover as yet unknown species where they occur. This means stepping up sampling efforts, particularly in the tropics where most of this undiscovered diversity is to be found. Our results confirm the existence of a regional peak in diversity at tropical latitudes, essentially explained by a strong beta component (Phillips et al., 2019; Goulpeau et al., 2024), while emphasising that it mirrors a considerable taxonomic deficit. Finally, once sampling coverage has been improved, new taxonomic approaches based on turbotaxonomy (i.e. the rapid description of species new to science allowed by automated procedures and the use of molecular tools; Butcher et al., 2012; Fernandez-Triana, 2022) and integrative taxonomy (i.e. the integrated use of multiple, independent lines of evidence—e.g. morphology, genetics, ecology, and behavior—for species delimitation and description; Dayrat, 2005) will be necessary to improve our ability to describe new species at a consistent rate (Padial et al., 2010; Fernandez-Triana, 2022).

In conclusion, we propose an updated estimate of the global number of earthworm species, as well as arguments for considering earthworms as a potentially hyperdiverse group of organisms, which has hitherto not been considered as such due to a strong taxonomic knowledge deficit (i.e the Linnean shortfall as defined in Lomolino, 2004). This is particularly remarkable when considering that earthworms are among the most widely publicised soil macro-organisms, due to their acknowledged importance in soil functioning. It is likely that this situation could also apply to other less well-studied soil taxa, leading to a considerable reassessment of the contribution of these organisms to global biodiversity.

## Acknowledgments and funding

The French Guiana dataset was acquired as part of different projects: DIADEMA (Dissecting Amazonian Diversity by Improving a Multiple Taxonomic Group Approach), DIAMOND (Dissecting and Monitoring Amazonian Diversity) and Wormbank (DNA barcoding earthworms in biodiversity hot spots of French Guiana) projects funded by “Investissement d’Avenir” grants managed by the Agence Nationale de la Recherche (CEBA: ANR-10-LABX-25-01; TULIP: ANR-10-LABX-41); “Our Planet Reviewed” French Guiana 2015 expedition (Touroult et al., 2018) organised by the Muséum national d’Histoire naturelle (MNHN, Paris) and Pro-Natura international in collaboration with the Amazonian Park of French Guiana, and financed by the European Fund for Regional Development (FEDER), the Regional Council of French Guiana, the General Council of French Guiana, the Direction de l’Environnement, de l’Aménagement et du Logement and by the Ministère de l’Éducation nationale, de l’Enseignement supérieur et de la Recherche. Additional support was received from the Réserve naturelle des Nouragues (Nouragues grants 2010 and 2011), the Réserve naturelle de la Trinité and the Parc Amazonien de Guyane.

The dataset from temperate France was obtained through a project called GloWorms, mostly funded by the ‘AgroEcoSystems’ Scientific Department of INRAE (National Institute for Research in Agriculture, Environment and Food, Paris, France). Samplings in Normandy were funded by the INPN (National Inventory of Natural Heritage, Paris, France). Sylvain Gérard’s PhD thesis was funded by a thesis grant from the Ecole Normale Supérieure (Paris, France). Daniel F. Marchán’s contribution was funded by the Make Our Planet Great Again programme (Campus France, Paris), via a postdoctoral grant.

JM and GGB acknowledge the FaunaServices group jointly funded by the synthesis center CESAB of the French Foundation for Research on Biodiversity (FRB; www.fondationbiodiversite.fr)”, CEBA, Sinbiose, Fapesp and CNPq (Brazil). NE gratefully acknowledges the support of iDiv, which is funded by the German Research Foundation (DFG – FZT 118, 202548816), as well as by the DFG (Ei 862/29-1, Ei 862/31-1). HRPP was funded by the European Union’s Horizon 2020 research and innovation program under the Marie Skłodowska-Curie grant agreement No. 101033214 (GloSoilBio). GGB acknowledges project support by the Brazilian National Council for Scientific and Technological Development (CNPq Nos. 312824/2022-0, 441930/2020-4 and 404191/2019-3).

## Conflict of interest disclosure

The authors declare that they comply with the PCI rule of having no financial conflicts of interest in relation to the content of the article.

## Data, scripts, code, and supplementary information availability

Data and scripts are available online on GitHub: https://github.com/EricMarcon/A_Can_Of_Worms-Appendix.

## Notes

https://github.com/EricMarcon/A_Can_Of_Worms-Appendix

### Competing Interest Statement

The authors have declared no competing interest.

### Summary of Updates

The preprint has been reformated according to a PCI template.

